# Toward Large-Scale Numerical Modeling of the Cardiovascular System with up to 34 Billion Vessels

**DOI:** 10.64898/2026.05.22.727287

**Authors:** Wayne Newhauser, Maxwell Cole, Patrick Diehl, Juana Moreno, Hartmut Kaiser, R. Tohid, Noujoud Nader, Jeffery Chancellor

**Affiliations:** Department of Physics, Medical Physics and Health Physics Program, Louisiana State University, LA, USA; Mary Bird Perkins Cancer Center, LA, USA; Center for Computation & Technology, Louisiana State University, LA, USA; Applied Computer Science, Los Alamos National Laboratory, NM, USA; Department of Computer Science, Louisiana State University, LA, USA; Department of Medical Physiology, Texas A&M University, TX, USA

**Keywords:** Cardiovascular, Computational model, Whole organism, Biophysics, High-performance computing

## Abstract

Cardiovascular diseases, such as stroke and heart attacks, are the leading cause of death worldwide. Computational models like cardiovascular digital twins (CVDTs) offer a promising path for research and intervention but are challenged by the complexity of simulating the full human vasculature. This study evaluates the feasibility of simulating blood flow through a vascular network containing 34 billion vessels (the estimated number in the human body) using first-principles physics and simplified geometry which is a first step towards CVDT.

We synthesized 3D vasculature using a fractal model and computed blood flow rates via Poiseuille equation and steady-state fluid dynamics, implemented with high-performance computing. Simulations were conducted for networks ranging from 6 vessels to 34 billion vessels.

The results demonstrated high accuracy (within 1% of benchmarks), reproducibility across platforms, and strong scalability. Simulating the full vasculature required 156 node-hours on the second-fastest supercomputer in the world, using 29 TB of memory and 84 TFLOPS. Maximum speedup factor was 80, with parallel efficiency no lower than 0.48.

These findings show it is computationally feasible to simulate blood flow through a full-body vascular network at scale. The approach is well suited to parallel computing, suggesting that with continued development, CVDTs could enable whole-organism modeling for applications such as stroke, trauma, radiation injury, and cancer metastasis.

## Introduction

Cardiovascular disease (CVD) has proven refractory to a wide range of management strategies, including varying degrees of socialized medicine to expand access to care (1), therapeutic drugs, surgical procedures, implanted devices, and behavioral modification. The high cost of CVD care delivery is a major challenge, *e*.*g*., in 2023 the US expended approximately a half trillion USD to reducing the burden of CVD (2, 3), corresponding to approximately 2% of its gross domestic product (4). In theory, expanding access to the current standard-of-care for CVD could provide substantial health benefits to some recipients. In practice, however, its high cost precludes a broad expansion (2). Hence, it is imperative to open new avenues of inquiry to find new standards of care that reduce costs and improve health outcomes. In 2019, the total cost of CVD in the US was over 400 billion USD, with 60% coming from direct health care expenses (5). In 2023, CVD research comprised only about 7% (4 billion USD) of total research funding allocated by the National Institutes of Health (NIH) (6, 7). Together, these issues comprise a strong economic impetus for research breakthroughs that will reduce the cost of CVD.

How might these future breakthroughs be achieved? Perhaps by application of traditional methods, including animal model experiments, human clinical trials, epidemiological studies, and others. These approaches certainly have increased our understanding of CVD and improved outcomes for many patients. Yet, despite decades of research progress accomplished with these methods, they have severe limitations, as evidenced by the fact that one third of all deaths worldwide are still caused by CVD (8). Hence, there is a strong public-health case to instigate non-traditional CVD research methods, especially those with the potential to transform CVD care.

One such approach is computational modeling, which is advancing rapidly. Explosive growth in computing power is enabling this approach. Anticipated future increases in computation speed, data transmission speeds, and data storage capacity will enable researchers to develop vascular models that are realistic, fast, and highly capable. In fact, it is theoretically possible to simulate some or all the key biophysical features and functions of the cardiovascular system of an individual, a type of model commonly called a ‘digital twin’. Cardiovascular digital twins (CVDT) will likely be useful for basic, applied, and clinical research, as well as personalized health care. These tantalizing prospects prompt critical questions, namely: ‘Will a CVDT, even in simplified form, become computationally feasible within years or decades?’, ‘If so, what scale of computing resource will be needed?’, and ‘Are the basic biophysical algorithms of CVDTs scalable using parallel computing systems?’ In this study, we sought answers to these open questions and, more generally, to guide future CVDT research, particularly with respect to scalability.

In stark contrast to knowledge of scalability, much is known about relevant computational research methods that could be brought to bear on cardiovascular digital twins (CVDT). The current capabilities of computational vascular research methods, *e*.*g*., involving one or a few vessels, are expansive and growing rapidly. These are built on solid foundations of biological and physical mechanisms. And much progress has been made in the application of high performance computing (HPC) to expand the capabilities of various vascular models (9). Taken together, these suggest that CVDTs could revolutionize vascular research, just as general-purpose Monte Carlo simulation codes, such as MC-NPX (10) and GEANT (11), transformed research in nuclear medicine, radiation therapy, diagnostic radiology, and other radiation sciences. The Monte Carlo method was invented in the 1940s, but its widespread translation into medical specialties did not occur until the 2000s, after sufficient computing resources had become available. Similarly, once computing resources become sufficient, a general-purpose vascular simulation code is expected to lead to new discoveries, facilitate experimental designs, test hypotheses, and CVDTs for clinical applications. Examples include intravenous drug delivery (*e*.*g*., to assist in the interpretation of cardiac images from nuclear medicine procedures, to diagnose traumatic brain injury, and to reduce the risk of radiation necrosis in cancer patients.

The theoretical bases already exist to develop a digital twin based on first principles, *e*.*g*., foundational theories regarding relevant aspects of physics (*e*.*g*., fluid dynamics, molecular diffusion, and radiation transport), biology and medicine (*e*.*g*., vascular anatomy, physiology, and hematology), and computer science (*e*.*g*., parallel computing, numerical methods). Many practical obstacles remain, however, including limitations in modeling capabilities, such as the number of vessels, their anatomical structure, the flow of blood, biophysical responses to injury and other stimuli, and time dependencies. These obstacles are caused in part by the sheer size of the problem. For example, tissue contains approximately 600 blood vessels per cubic millimeter (12), or 34 billion vessels in the entire human body, corresponding to more than 150, 000 km in length (13). The complete vasculature is so massive that it was considered computationally infeasible. Several recent studies, summarized below, reported breakthroughs that, taken together, suggest a first digital twin of the human vasculature may become computationally feasible within the next few years. Evidence of computational feasibility is a crucial milestone on the path toward a general-purpose vascular simulation code. The need for such evidence comprises the impetus for this study.

Modeling approaches may be categorized as bottom-up or top-down. These complementary approaches are both necessary to develop CVDTs. Most computational vascular research is bottom up, *i*.*e*., with special-purpose codes that omit or approximate many CV features. For example, by omitting most blood vessels, it becomes feasible to model small numbers of vessels with remarkable detail in terms of vessel geometry, fluid dynamics, and other phenomena. Linninger *et al*. citelinninger2019mathematical combined image-based mesh modeling with a stochastic space-filling algorithm to model the vasculature of a mouse brain hemisphere with a high degree of anatomical accuracy, including nearly 500, 000 vessels. For context, the human brain contains about 9 billion vessels (14, 15). In a contrasting approach, our laboratory has focused on modeling remarkable numbers of vessels, where vascular geometry and fluid dynamics were simplified. For example, Donahue *et al*. (16) simulated 17 billion vessels using a simple fractal vascular network. Table 1 summarizes the current state of the art in digitally modeling large numbers of vessels.

**Table 1.** Selected computational vascular models in the literature.

| Organ | Species | Number of Vessels | Source |
| --- | --- | --- | --- |
| Cerebral arterial tree | Human | 300 | (17) |
| Retina | Mouse | 1,250 | (18) |
| - | - | 8,000 | (19) |
| Heart | Canine | 43,000 | (20) |
| Subsection of cerebral cortex | Human | 256,000 | (21) |
| Brain hemisphere | Mouse | 500,000 | (22) |
| Fat pad | Mouse | 1.7 million | (23) |
| Brain | Human | 9-17 billion | (16, 24) |

The quantity blood flow can be modeled with simplifications to achieve computational feasibility. The most complete and accurate approach is to solve the three-dimensional, time-dependent Navier-Stokes equations. For example, Xiao *et al*. (25) modeled arterial networks using this method in networks containing less than 50 vessels. Updegrove *et al*. (9) developed the SimVascular code to create vascular networks from volumetric images and to calculate blood flow in the body’s major arteries. A special case of the Navier-Stokes equations, known as Poiseuille’s equation, applies to the simpler scenario of steady-state flow of an incompressible Newtonian fluid in a cylindrical pipe of constant cross section. Using this simplification, Donahue *et al*. (16, 24) modeled blood flow in a vascular network with up to 17 billion vessels using high-performance computing methods. These examples illuminate some of the tradeoffs between modeling realism and computational expense. Together, they strongly suggest it will become feasible to develop a general-purpose code, which would model all the vessels in the human body with sufficient detail and speed for several types of vascular research, including those involving radiation exposure. This could comprise a hybrid model (Fig. 1), in which diverse computational vascular models are combined to create a whole whole-body model. How to accomplish this is poorly understood and will involve considerations of achievable scale, realism, computation time, and computing systems.

**Fig. 1.**
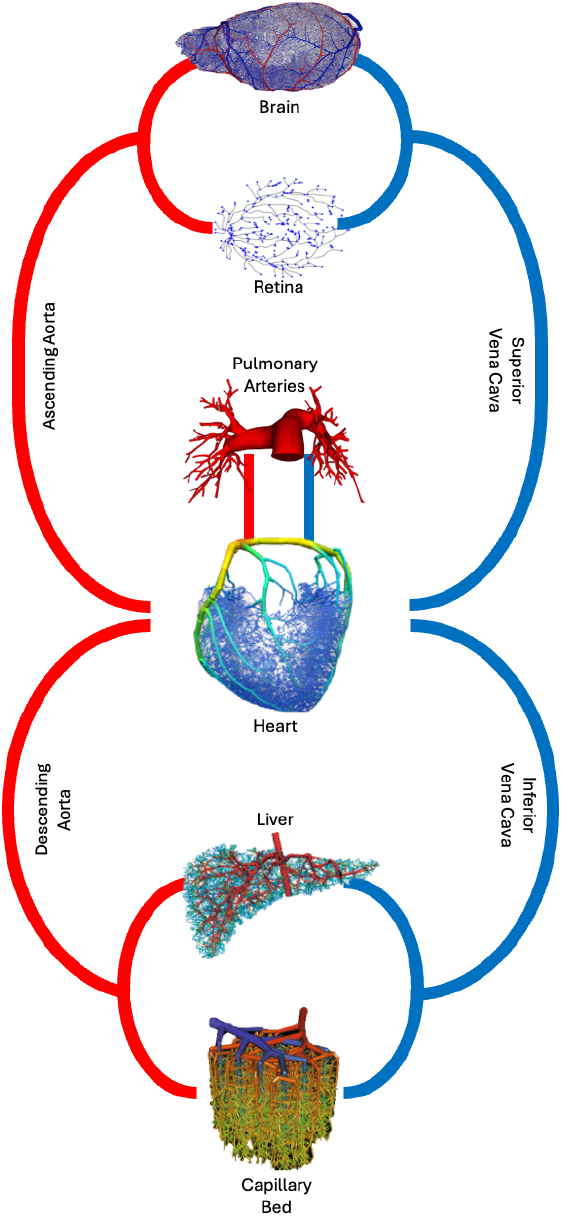
Hybrid computational vascular model. In this illustration the brain model is taken from Linninger *et. al*. (22), the retinal model from our laboratory (Nader N., Newhauser W., Cole M., Diehl P., Tohid R., and Kaiser H.), the pulmonary arteries from SimVascular (26), the heart model from Hyde *et. al*. (20), the liver from Barleon *et. al*. (27), and the capillary bed from Linninger *et. al*. (21).

It appears that a full-scale CVDT requires vastly more memory and computing operations than are available from a single computer. Parallel computing systems overcome these limitations by virtue of their inherent ability to scale, namely, to accommodate increasing amounts of data and workload. Little attention has been paid to scaling rules for CVDTs. These are needed to inform future CVDT computational study designs, *e*.*g*., to estimate the amount of time required to simulate blood flow through a specified number of vessels, given a specified level of biophysical detail. Donahue *et al*. (24) proposed scaling rules for networks of up to 17 billion vessels, radiation injury and consequent changes in blood flow. To our knowledge, no other scaling rules have been reported for CV modeling. The previous literature contained no reports on the impacts of anatomic realism or the architecture of high-performance computers on scaling rules.

The primary objective of this study is to demonstrate the feasibility of simulating blood flow through a vasculature of the order of the human body, or 34 billion vessels. Secondary objectives of this study were to develop scaling rules to estimate the achievable performance envelope, *e*.*g*., the speed, scalability, and cost of various types of cardiovascular calculations. We used several high-performance computing clusters of diverse size and architecture, including the world’s second fastest supercomputer to assess portability, scaling, and reproducibility of a prototype CVDT.

This work represents the first prototype (P1) of a foundational base model on the path toward a personalized vascular digital twin. The overall development strategy and projected timeline are illustrated in Figure 2. Prototype 1 establishes the computational framework and demonstrates large-scale feasibility: for the first time, we show that it is computationally possible to simulate the full vascular network of approximately 34 billion vessels using a simplified geometric representation. The purpose of this prototype is not physiological completeness, but rather to validate scalability, numerical robustness, and architectural design at whole-body scale. Building upon this foundation, Prototype 2 (P2) extends the model to incorporate pulsatile flow dynamics (publication in preparation), thereby introducing physiologically relevant time-dependent hemodynamics. Prototype 3 (P3) further advances the framework by modeling vessel rupture at the level of individual vessels using a nonlocal bond-based formulation (28). This step introduces structural failure mechanisms and enables the study of pathological scenarios. Finally, Prototype 4 (P4) will incorporate anatomically accurate vessel geometries and organ-specific structures, significantly increasing physiological realism and enabling subject-specific adaptation. Together, these prototypes form a structured and incremental road map toward a personalized digital twin. Each stage systematically increases physiological fidelity while preserving computational scalability, ensuring that the final framework remains both patient-specific and computationally tractable.

**Fig. 2.**
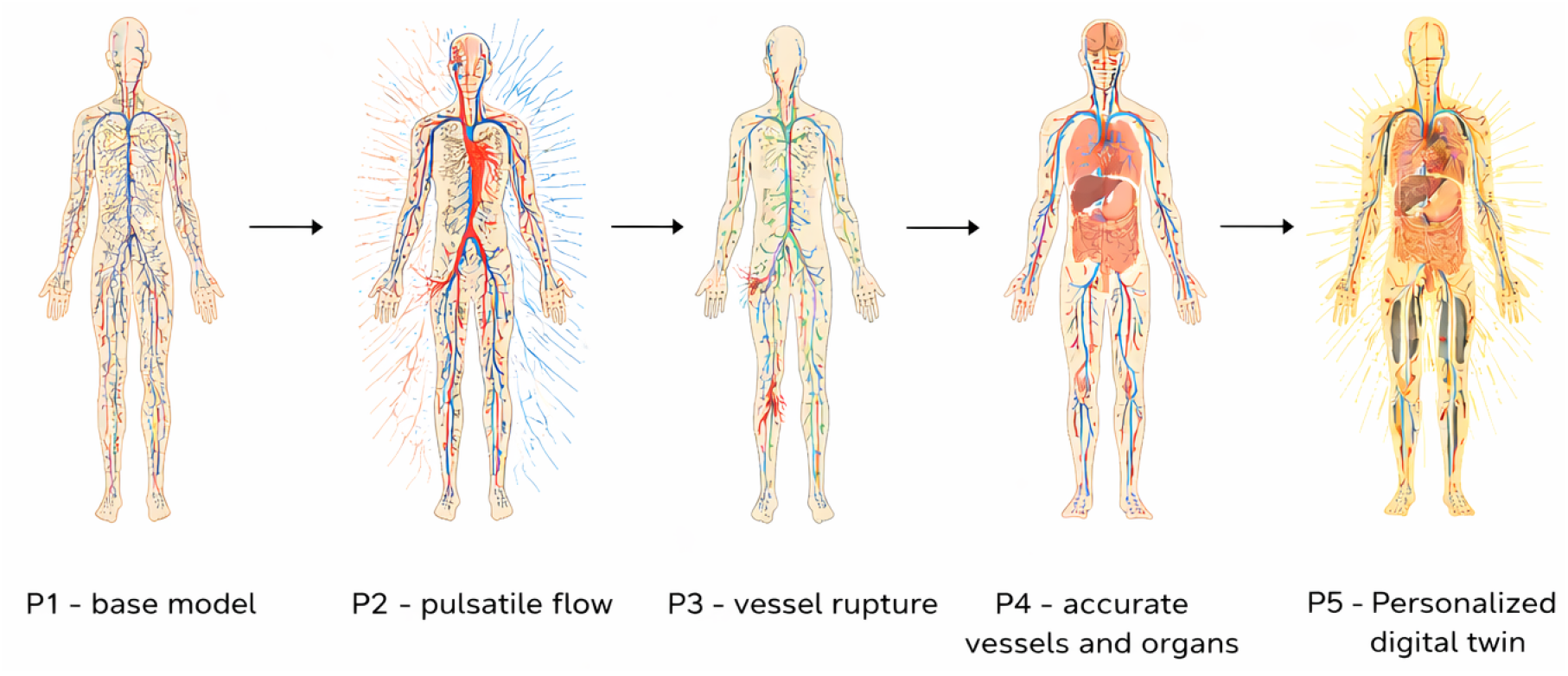
Time line towards the personalized digital twin: P1 – computational model with 34 billion vessels (this work); P2 – adding pulsatile flow to this work; P3 – model vessel rupture using peridynamic modeling; and P4 – adding more accurate vessel geometry and modeling organs.

## Methods

We built a three dimensional computational model of the human vascular system by creating computer codes to generate vessels, pre-process the network of vessels into optimized matrix form, and solve matrices to yield blood pressures and blood flow rates throughout the vascular network. The blood vessels were generated using a fractal model with right-left mirror symmetry of the arterial and venous trees. The next step, called preprocessing, partitioned the vascular network and communicated partition information across compute nodes in a cluster of parallel computers. The final step utilized a vessel-conductance matrix to solve for blood pressure values at each node to calculate blood flow rate through each vessel. Flow rates were calculated using Poiseuille’s equation, a special case of the more general Navier-Stokes equation of fluid dynamics.

The approaches in this work follow those of Donahue *et al*. (16, 24). All computer codes were developed anew for this study to further increase their capacities, including the number of vessels considered, speed, portability, ease of use, reliability, and modularity. To accomplish these, we simplified the software architecture, reduced the number of third-party libraries used, eliminated the use of proprietary libraries, and exclusively utilized open-source compilers and libraries. Several algorithms and numerical methods we used earlier (16, 24) were replaced with functionally equivalent alternatives. We retained several validation methods and data from our previous studies to facilitate comparisons with the earlier results. These included benchmarking tests, validation calculations based on computational fluid dynamics (CFD), and scalability metrics to characterize performance.

We developed two versions of each code. A low-performance version of the code was written in the Python language to facilitate development and testing using small-scale simulations. A high-performance version was written in the C++ programming language for use on large-scale simulations running on clusters of parallel computers.

## A. Overview of Algorithms

### A.1. Vascularization

We synthesized networks of arbitrary number of vessels in three dimensions for blood-flow simulations and, to facilitate visualization, in two dimensions. The approach used a fractal geometry (16, 29, 30) to generate an arterial tree that fed a mirror-symmetric venous tree (Fig. 3). The fractal geometry was essential because it is the only known strategy capable of synthesizing the number of vessels in the human body. The vessels were modeled as straight, rigid conduits. In a tree, each parent vessel bifurcated into two smaller daughter vessels at a junction. Each stratum of daughter vessels was called a generation. This process repeated until the desired number of generations of vessels was reached. The total number of vessels in a network, *N*_*V*_ , was specified by selecting the number of generations, or

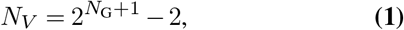

where *N*_G_ is the number of generations. In the final generation, the daughter arterial vessels were joined to the corresponding venous ones at the midplane of the vasculature

**Fig. 3.**
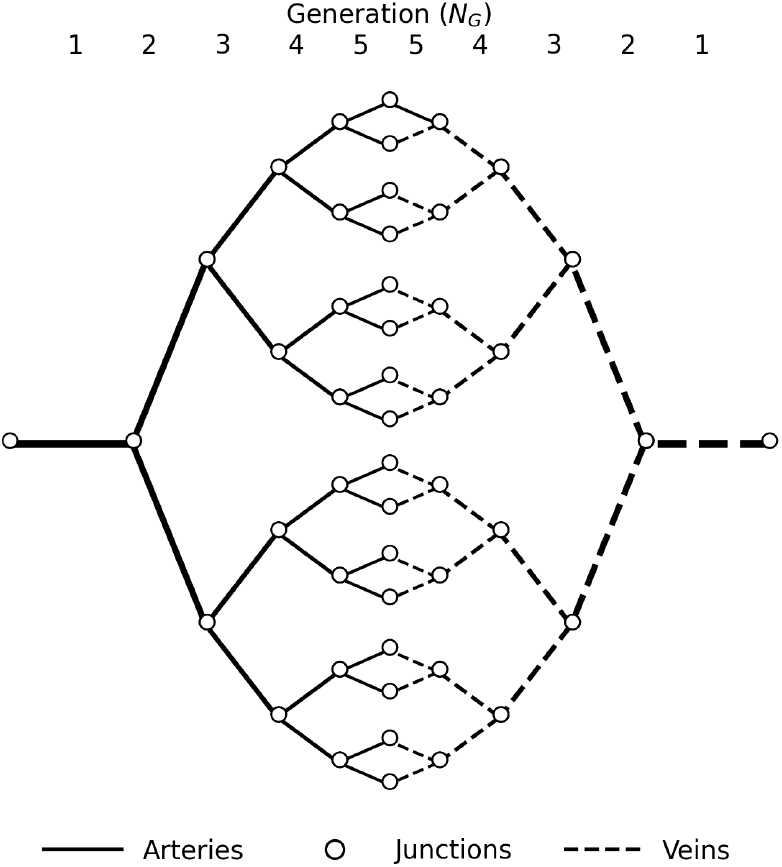
An example of a 2-dimensional 5-generation fractal vascular network in the x-y plane. The solid lines represent arteries, and the dashed lines represent veins. The white circles denote vessel junctions.

The fractal model was configured to provide biologic realism in terms of the number of vessels in each generation and the vessel density. Model parameters included the number of generations (*N*_G_), the length of the smallest vessel (*L*_min_), the radius of the smallest vessel (*r*_min_), and the initial angle of bifurcation (Θ_0_). We defined the reduction factors for length (*ξ*_*L*_), radius (*ξ*_*r*_), and angle of bifurcation (*ξ*_Θ_), to correspond with reference anatomical data. Table 2 lists the parameter values.

**Table 2.** Parameters for generation of the vascular network.

| Parameter (units) | Symbol | Value | Source |
| --- | --- | --- | --- |
| Number of generations | $N_G$ | $2 \leq N_G < 34$ | This work |
| Length of smallest vessel ($\mu\text{m}$) | $L_{\min}$ | $55 \mu\text{m}$ | (31) |
| Radius of smallest vessel ($\mu\text{m}$) | $r_{\min}$ | $2.5 \mu\text{m}$ | (32) |
| Initial bifurcation angle ($^\circ$) | $\Theta_0$ | $37.47^\circ$ | (33) |
| Vessel length scaling factor (rel. units) | $\xi_L$ | 0.78 | (16) |
| Vessel radius scaling factor (rel. units) | $\xi_r$ | $2^{-1/3}$ | (34) |
| Bifurcation angle scaling factor (rel. units) | $\xi_\Theta$ | 0.88 | (35) |

The fractal algorithm began by determining the geometrical extents of the vascular network, *i*.*e*., the starting x-coordinate of the arterial tree, the ending x-coordinate of the venous tree, and the maximum or minimum y-z bounds in of the final generation. We synthesized vessel trees beginning from the outside and worked inward toward the midplane, where the two trees meet. At the end of each vessel, a junction was established, where the vessel then bifurcated into two daughter vessels, as shown in Fig. 4.1 and 4.2. The pair of symmetric daughter vessels was synthesized with reduced radius, length, and angle of bifurcation using the reduction factors in Table 2. To calculate the ending coordinate of each vessel, we extended from its starting junction with a length of 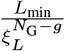, where *g* is the current generation number of the vessel, and with angle 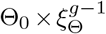. The vessel radius is calculated by 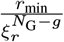. The synthesis process was repeated until *N*_G_ generations were reached, at which time the vessels of arterial and venous trees met at the midplane. For 3-dimensional networks, generations of bifurcated vessels were alternately created in orthogonal *x*-*z* and *y*-*z* planes.

**Fig. 4.**
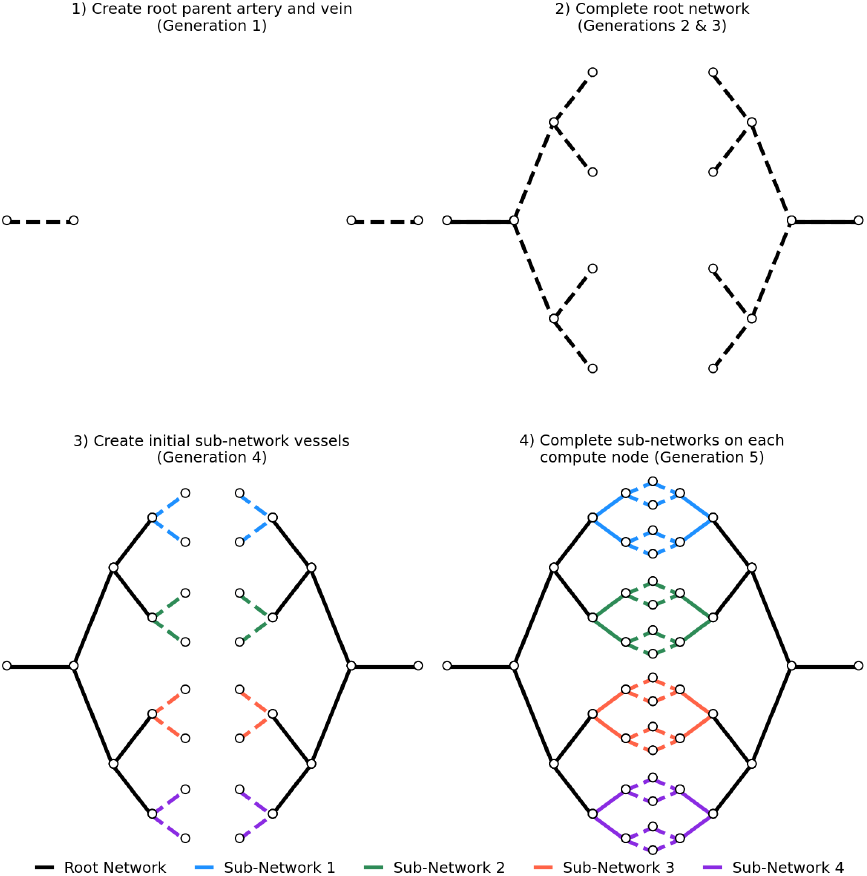
Sequence of steps for distributed creation of a 5-generation network on 4 compute nodes. The dashed lines represent newly created vessels in a given step.

The algorithm stores a set of vessel descriptors, which include each vessel’s start and end coordinates, radius, and indices of connected junctions. With this approach, each vessel descriptor requires just 72 bytes. A complete set of descriptors (for 34 billion vessels of the whole body) required nearly three terabytes of memory, more than is available on a single compute node in a typical high-performance computer. To overcome this memory limitation, we distributed the vessel synthesis by dividing the vessel network into an arterial root tree, a venous root tree, and multiple subnetworks that connected to the root trees (Fig. 4). Each compute node created both root trees and one unique subnetwork. We included the root trees on each compute node because they contain needed information on the start and end junctions of its subnetwork. The size of root trees and subnetworks varies with the number of generations, *N*_G_. The number of generations in a root tree is

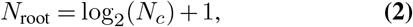

where *N*_*c*_ is the number of compute nodes (16). The number of generations within a subnetwork is *N*_G_−*N*_root_ (Fig. 4). Sub-networks were synthesized in parallel using the Message Passing Interface (MPI) (36).

### A.2. Partitioning & Preprocessing of the Vascular Network

We utilized the method of distributed-memory parallelization using MPI to assemble the vascular network across multiple compute nodes. Because the vascularization algorithm (Section A.1) stored vessel descriptors in order of generation, many junctions were distributed to, or shared by, several compute nodes. These shared junctions, sometimes called edge junctions, increased inter-processor communications and thereby strongly increased the execution time. Therefore, we developed a partitioning algorithm to minimize the number of edge junctions and attendant inter-process communication. This, in essence, was accomplished by allocating only contiguous groups of vessels to any given compute node (Fig. 4). In this algorithm, a “head compute node” was selected to store the root network data, while all remaining compute nodes stored the data only from their assigned sub-networks. Thus, the partitioning algorithm limited the number of shared junctions to be equal to the total number of compute nodes used. Partitioning also provided static load balancing by assigning approximately the same number of vessels to each compute node.

Prior to the commencement of blood flow calculations, the partitioned vascular network data was preprocessed. In this step, the head compute node identified each pair of adjacent subnetworks with shared junctions (see Fig. 4) and communicated this to each of the respective pairs of compute nodes. The original indices of the vessel junctions were then renumbered to be consistent with the new partitioning scheme.

### A.3. Blood Flow Calculations

The Navier-Stokes equations are commonly used in computational fluid dynamics for complex blood flow analyses (37). These equations can be simplified for laminar flow in a pipe of uniform cross-section, with some assumptions: the fluid flow is steady, axisymmetric, fully developed, and there are no radial or azimuthal velocity components. The volumetric flow rate, *Q*, in each vessel can then be described by a specialized case of the Navier-Stokes equation, known as the Poiseuille equation, or

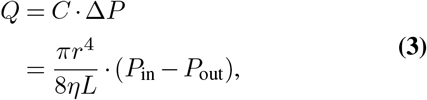

where *C* is the vessel conductance, *r* is the vessel inner radius, *L* is the vessel length, *η* is the blood viscosity, *P*_in_ and *P*_out_ are the blood pressures at the network inlet and outlet, respectively (38). The Poiseuille equation gives the pressure difference of an incompressible Newtonian fluid in a cylinder. The blood viscosity was taken from an empirical formula from Pries *et al*. (39) for a constant hematocrit of 0.45, or

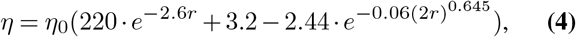

where *r* is the vessel inner radius in millimeters, and the value of *η*_0_ is 35 mPas. This viscosity model was selected because it can be evaluated without *a priori* knowledge of hemodynamic values and because it has been previously bench-marked for microvascular networks. As a simplifying assumption, we treated the blood as incompressible and the vessel walls as impermeable to fluids. Therefore, the net fluid flow rate at each junction was zero (40). The flow rates at each junction of the network could, therefore, be cast as a system of linear equations. The system of equations for a 6-vessel network was

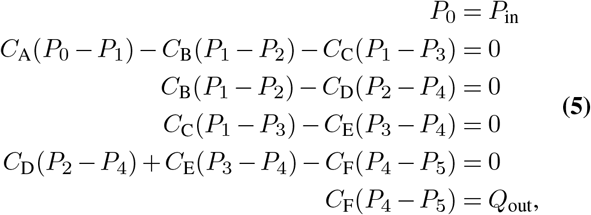

where *P*_in_ is the inlet pressure and *Q*_out_ is the outlet blood flow rate. We took the inlet boundary condition (*P*_in_) to be the mean output blood pressure of the heart in the average resting adult, 100 mm Hg (13.3 kPa) (41). The outlet boundary condition (*Q*_out_) was the mean blood flow of 26 ml/s for a resting adult as reported by Joseph *et al*. (42)

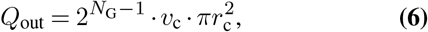

where *N*_G_ is the number of generations in the network, *v*_c_ is the blood velocity in the smallest vessel, and *r*_c_ is the radius of the smallest vessel (24). For *v*_c_, we assigned a value of 1 mms^*−*1^ taken from Ivanov *et al*. (43). This system of equations was put in matrix form, see Eq. 7.

The 2-dimensional matrix at left, called the conductance matrix, described the conductance value of each vessel. The 1-dimensional pressure vector described the blood pressure value at each junction, and the flow vector on the right hand side described the net flow rate at each junction or the boundary conditions. Each compute node was assigned a partition from the Partitioning & Preprocessing algorithm (Section A.2) and constructed those rows of the conductance matrix (in compressed sparse row format) for each junction that it was assigned.

The matrix (Eq. 7) was solved to obtain the unknown blood pressure values at junctions. The pressure values for all shared junctions were then communicated to the corresponding compute nodes. Pairs of inlet and outlet pressure values were then substituted into (Eq. 3) to calculate the blood flow rate through each vessel. The flow rates were written to persistent storage for analysis.

## B. Computational Implementation

### B.1. Software and Hardware Environment

A major challenge in large-scale computational CVD research is the diversity of computer architectures available to researchers, *e*.*g*., processors from different manufacturers. We designed our codes to run, without modification, on architectures ranging from a laptop computer to supercomputers.

Two supercomputers provided data on the trade-off of speed and cost. Supercomputer Fugaku (44), the second fastest supercomputer in world at the time of this study (now fourth fastest), with maximum double-precision performance of 442 PFLOP/s (45). Supercomputer Fugaku contained 158, 976 1.8-GHz compute nodes. Each compute node had one 48-core processor (ARM A64FX), with a double-precision performance of 2.76 TFLOP/s, 32 GB of shared memory, and an energy efficiency of 54 GFLOP/W. Perlmutter (46) was the 7^th^ fastest supercomputer in the world at the time of this study (now 14^th^ fastest) at 70 PFLOP/s (45). Perlmutter contained 3072 424-GHz compute nodes, each with two 64-core processors (*AMD EPYC 7763*), 512 GB of memory, and an energy efficiency of 27 GFLOP/W.

We compiled all codes, written in the C++ programming language, using a freely available compiler (the GNU Compiler Collection 12) running on open-source operating systems (Linux kernel 4.18). Distributed memory parallelization was utilized through an open-source message passing interface (OpenMPI 4.1) (36). All output data was written to permanent storage files in binary format. We used linear algebra algorithms from the Trilinos Project (47), including matrix objects from the `Tpetra` package (48), preconditioners from `Ifpack2` (49), and the parallel matrix solver from `Belos` (50). The conductance matrix was solved using the generalized minimal residual (GMRES) iterative Krylov method (51). To increase the rate of convergence, we pre-conditioned the matrix using incomplete lower-upper (ILU) factorization with a fill-level of 2. The GMRES method for all networks required fewer than 100 iterations to reach a convergence tolerance below 1 ×10^*−*12^. We also used a linear algebra template library (Eigen v3.4.0) for optimized sparse matrix linear algebra routines (52).

### B.2. Performance Characteristics

By virtue of its vast size and complexity, the complete cardiovascular system is difficult and time consuming to simulate. Indeed, simulations that include all 34 billion vessels and high realism will require much more computing resources than are currently available. The anticipated growth in computing power and limitations of performance improvement have been studied for decades, *e*.*g*., Moore’s Law, Amdahl’s Law (53–55). In contrast, the literature is sparse on computational scalability in CVD simulations (16, 22, 24), and previous attempts to understand scaling laws in this domain were almost exclusively based on observations that are poorly suited for extrapolation to larger scales. As a result, it is nearly impossible to ascertain with any degree confidence what type of CVDT simulations will become possible in the near future, much less in the long term. For these reasons, to design future research studies, there is an urgent need for tools to quantify and assess how much computing power CVDT simulations will consume and how they will scale using parallel computing methods.

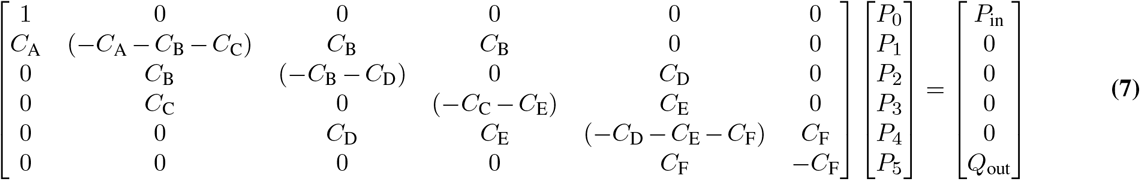

To characterize the scalability of vascular simulations, we implemented our codes on two high-performance computing clusters (see Section B.1 for details). We measured simulation times and ascertained the maximum number of vessels that can be simulated on a given cluster by iteratively increasing the size of the vascular network and recording the simulation times. Simulations ranged from 6 vessels to 34 billion vessels.

Simulation time depended strongly on both features of the cardiovascular model the computing architecture that were used. These features are conceptually distinct but inextricably linked. We assessed them both by measuring simulation times, as well as metrics that are independent of computing architecture, *e*.g., the total number of floating-point operations required, scalability, and electrical energy usage. With these measured data, we developed scaling formulae, described below, that relate network size to computing time and economic cost. The formulae facilitated the design of subsequent computational experiments in this work.

When designing large computational experiments, it is commonly necessary to first estimate the computing time that will be required. We estimated the minimum amount of wall-clock time required as

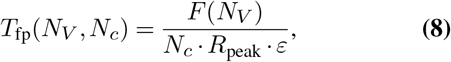

where *F* (*N*_*V*_) is the number of floating-point operations required as a function of number of vessels, *N*_*V*_ , and *R*_peak_ is the maximum rate at which a processor computes floating-point operations (45). More precisely, *R*_peak_ is a theoretical maximum computation rate of the CPU that purposefully excludes performance losses caused by inefficiencies in users’ codes. *ε* is the parallel efficiency of the software running in parallel on the cluster’s nodes, and *N*_*c*_ is the number of compute nodes used. Each of these terms are described in more detail below.

Increasing the number of compute nodes can yield substantial performance improvements. However, at some point, adding more compute nodes yields diminishing returns. Scalability metrics attempt to characterize the ability of the hardware and software to utilize increased computational resources. Such increases typically refer to additional compute nodes, which provide greater computing power and more electronic memory. The number of compute nodes and the amount of memory impact scalability in different ways and are, therefore, estimated with distinct methods. Specifically, we used the quantity speedup factor, *S*, to assess the scalability of the simulations when bound by limitations in compute-node performance (*i*.*e*., strong scalability). We used the quantity of parallel efficiency, *ε*, to assess the (weak) scalability of the simulations in cases bound by limitations on the amount of electronic memory available on individual compute nodes. An important practical application of *S* is to estimate the time required for a fixed problem size, with the goal of reducing wall clock time required. Similarly, *ε* was used to estimate the time required for increasingly large problem sizes, with the goal of maintaining approximately constant wall clock time. *S* and *ε* depend on the problem size and how that quantity is defined.

In this study, we defined the problem size in a manner to mainly include enduring physical aspects of the vascular model (*e*.g., the number of vessels in the human body). We also sought to mostly exclude aspects that will soon be rendered obsolete by rapid advances in hardware technologies. To that end, we defined “problem siz” to include only the number of floating-point operations. We intentionally excluded integer computations, memory access requests, inter-process communications, and input/output operations, among other forms of computational overhead that are highly dependent on hardware. We used a profiling tool (Linux perf tool (56)) to measure the sum of floating-point operations, and fit the values to an empirical approximation given by

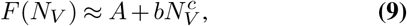

where *A* is a code-specific constant that reflects the program’s computational overhead, and 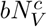 is a power law representing the dependence on the number of vessels simulated.

The ability to utilize additional resources for a fixed problem size is described by strong scaling, measured by the speedup factor given by

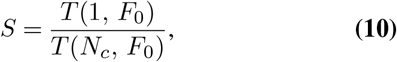

where *T* (1, *F*_0_) is the wall-clock time required to execute a task of problem size *F* = *F*_0_ using a single compute node, and *T* (*N*_*c*_, *F*_0_) is the wall-clock time required to perform the same task using *N*_*c*_ compute nodes. A theory of strong scaling is embodied by Amdahl’s Law (54), where the speedup factor is given by

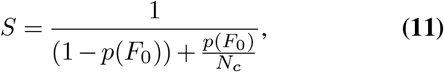

where *p*(*F*_0_) represents the fraction of execution time *T* (1, *F*_0_) that is spent on tasks that, theoretically, could have been completed faster if they had been computed in parallel (*N*_*c*_ *>* 1). Faster completion is possible because the additional compute nodes decrease the *workload-per-node*. As the number of nodes becomes large, the workload per node eventually becomes vanishingly small and the total execution time is predominated by serial tasks. In this case, the speedup factor approaches an asymptotic limit,

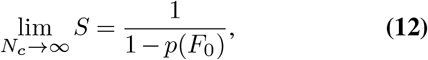

and further reductions in wall clock time become vanishingly small. Therefore, strong scaling and Amdahl’s Law are commonly used in cases where the goal is to reduce wall-clock time for a fixed problem size *F*_0_.

To evaluate Eq. (10) for *S*, we measured *T* (*N*_*c*_, *F*_0_) for vascularization, partitioning, and blood flow calculations on *N*_*c*_ = 1, 2, 4, 8, …, 1024. We chose a problem size of *F*_0_ = 23 GFLOP, corresponding to a 24-generation network (33 million vessels) because this was the largest network size achievable on a single compute node of the HPC systems used. We also fit measured *S* values from Eq. (10) to Amdahl’s Law, Eq. (11), to determine the value of *p*(*F*_0_). It is important to remember that strong scalability, in essence, characterizes a reduction in execution time when the number of nodes increases but the problem size is constant.

In contrast, weak scaling characterizes performance in situations where both the resources and problem size are increased. When the *problem size* increases proportionally with the number of compute nodes, the workload-per-node remains constant, and ideally the wall-clock execution time also remains constant. The quantity parallel efficiency characterizes the degree to which that ideal is achieved (57), or

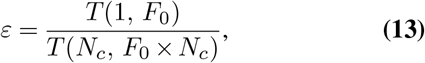

where *T* (*N*_*c*_, *F*_0_ × *N*_*c*_) is the wall-clock time for problem size *F* = *F*_0_ ×*N*_*c*_, *i*.*e*., the problem size is scaled in proportion to the number of compute nodes. A theory of parallel efficiency is embodied by Gustafson’s Weak Scaling Law (55), or

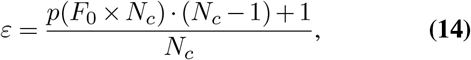

where *p*(*F*_0_ × *N*_*c*_) is the fraction of *T* (1, *F*_0_ × *N*_*c*_) that is spent on tasks that could be reduced if executed in parallel on multiple compute nodes. This theory reveals that, as the problem size and number of nodes increases, the parallel efficiency declines, eventually approaching the asymptotic limit given by

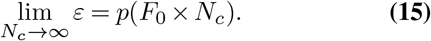

Parallel efficiency, from Eq. (13) and Eq. (14) is commonly used to estimate wall clock time, Eq. (8), in cases where the problem size and number of nodes vary together.

We determined *ε* from Eq. (13) using measured values of *T* (*N*_*c*_, *F*_0_ ×*N*_*c*_) for vascularization, partitioning, and blood flow calculations and *N*_*c*_ = 1, 2, 4, 8, …, 1024. The problem size *F*_0_ = 23 GFLOPs (*i*.*e*., *F*_0_ ×*N*_*c*_ = 23, 2 ×23, 4 ×23, …, 1024 ×23 GFLOP). We also fit measured *ε* values from Eq. (13) to Gustafson’s Law in Eq. (14) to determine the value of *p*(*F*_0_ ×*N*_*c*_), which is needed in Eq. (8) to estimate wall clock time.

As previously described, we analyzed the efficiency of our algorithms by assessing the rate of increase of the problem size, *F* (*N*_*V*_) from Eq. (9) as a function of the number of vessels, *N*_*V*_. In addition, to facilitate interpretation of this result, we also expressed it in Big 𝒪 notation (58, 59), which denotes the order of the algorithms. The order of our algorithms was then compared with a variety of commonly used algorithms for context. Based on the result of these comparisons, we assigned our algorithms to a qualitative category of efficiency. The rubric to assign qualitative categories was as follows: 𝒪 (1) or better was is considered high, between 𝒪 (1)and 𝒪 (*N*_*V*_) is moderate, and 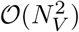 or worse is low.

The minimum electrical energy required, *E*, is empirically given by

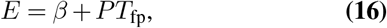

where *β* is a cluster-specific constant that represents the baseline amount of energy required to execute a simulation of any size, *P* is the electrical power consumption of a cluster’s compute node, and *T*_fp_ is the time required for floating-point operations. The power consumption rates were taken from previously measured values (45). We used open-source profiling tools (60) to measure the energy use of our algorithms. We then compared the measured values to those estimated by Eq. (16), where *T*_fp_ values were from Eq. (8).

The weak scaling, strong scaling, and energy consumption were characterized on two architecturally distinct HPC clusters (Section B.1).

### C. Validation

Validation methods follow those previously reported by our laboratory (16, 24) with modifications to accommodate the revised algorithms, the increased network size, and the different computational architecture used. Validation included tests for accuracy, reproducibility, and repeatability of the codes.

To validate that the values of conductance, blood pressure, and blood flow rate (Section A.3) were accurate, we compared them with previously validated regression-test data (16, 24). We additionally compared values of blood flow rates calculated with Eq. (3) to corresponding values calculated with a second method, described below. For a symmetric network with steady-state blood flow, in which the net flow at each junction is zero, it follows that the volumetric blood flow rate (*Q*) in each vessel is

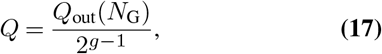

where *Q*_out_ is the outlet boundary condition flow rate, *N*_G_ is the number of generations in the system, and g is the generation number of the vessel. The blood flow rate values from Eq. (3) were considered valid if they agreed with the corresponding value from Eq. (17) within a relative difference of 0.05% for all vessels considered. We also validated the accuracy of predicted blood pressures from Eq. (7) and flow rates from Eq. (3) by comparing them to corresponding test results obtained using a previously validated commercial computational fluid dynamics simulation software (Autodesk CFD, San Rafael, CA, USA) (24) to ensure the maximum relative was less than 1% for all vessels considered. To ensure reproducibility, we implemented the algorithms in two programming languages, namely, C++ and Python, to test reproducibility on a personal computer (M1 processor with 8 GB of memory, Apple Inc., Cupertino, CA, USA). Additionally, we performed regression tests, as described above, using codes implemented in both the C++ and Python programming languages.

## Results

The cardiovascular digital twin codes exhibited scalability up to 34 billion vessels, as well as improved execution speed, portability, and reproducibility. The codes ran on multiple platforms without modification. Platforms considered include both shared memory (single CPU) and various distributed (computer clusters) environments. The 34-billion vessel CVDT required 84 TFLOP and 29 TB of memory. Results passed a battery of validation tests.

### D. Performance Characteristics

We measured the execution times of the vascularization, partitioning & preprocessing, and blood flow calculation algorithms for networks ranging between 128 vessels and 34 billion vessels, or 6 to 34 generations, using *N*_*c*_ = 1, 2, 4, 8, …, 1024 compute nodes. A single compute node could accommodate networks with up to 24 generations (33 million vessels), beyond which each additional generation required a factor of 2 increase in number of compute nodes.

Fig. 5 plots the total execution time (all inclusive, *i*.*e*., floating-point operations, input/output operations, latency, etc.) versus the number of vessels recorded on the Supercomputer Fugaku. For a network containing approximately the number of vessels in the entire human body (34 billion), the vascularization, partitioning & preprocessing, and blood flow calculation algorithms required a total of 156 node-hours (9 wall-clock minutes on 1, 024 compute nodes). For context, Fig. 5 also displays the sizes of a variety of vascular networks in biological specimen: approximately 100 arteries in the pulmonary system (61), 24, 000 vessels in the human retina (62), 500, 0004 vessels in the hemisphere of a mouse brain (22), 50 million capillaries in the human kidneys (63), 300 million vessels in the human lungs (64), 9 billion capillaries in the human brain (16), and over 34 billion vessels in the entire human body (12).

**Fig. 5.**
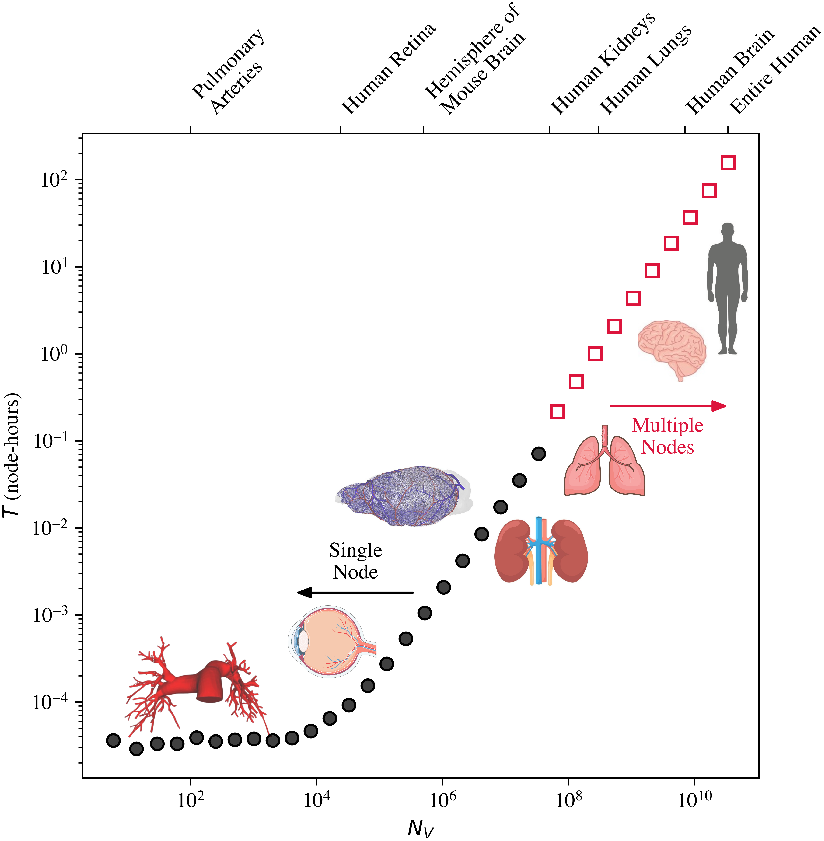
Plot of time, *T* , versus the number of vessels, *N*_*V*_ , required to execute all algorithms for networks ranging from 6 vessels to 34 billion vessels. The black circles represent runs that used a single compute node, and the red squares multiple nodes. The number of nodes used, for the red squares, double with each subsequent data point.

Fig. 6 plots the required number of floating-point operations for networks ranging in size from 6 vessels to 34 billion vessels. The measured values were fit to an empirical formula, Eq. (9), to allow for expedient estimations of floating-point operations for an arbitrary number of vessels. On the Supercomputer Fugaku, the parameters for Eq. (9) were determined to be *A* = 8.50 ×10^5^, *b* = 2.85 ×10^3^, and *c* = 0.994. On Perlmutter, the parameters were *A* = 1.70 ×10^4^, *b* = 2.58 ×10^3^, and *c* = 0.979. The differences in these values can be attributed to the compiler optimization schemes available on the utilized HPC clusters.

**Fig. 6.**
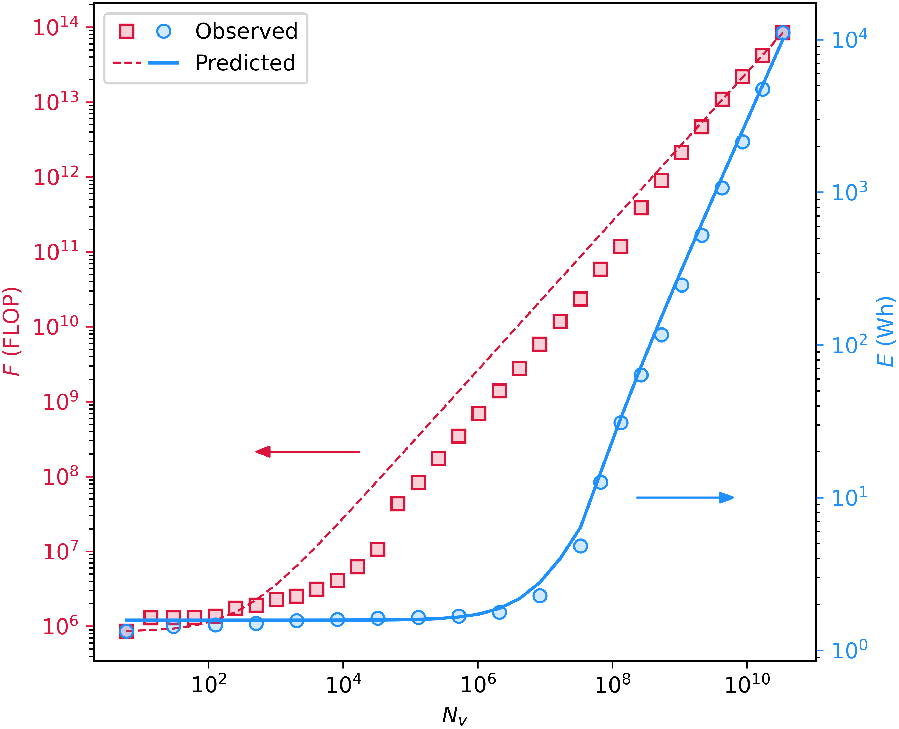
Floating-point operations, *F* (red, left axis), and energy, *E* (blue, right axis), required to simulate networks ranging between 6 and 34 billion vessels, *N*_*V*_. The red squares represent the measured number of floating-point operations and the dashed red line shows the empirical fit from Eq. (9). The circles represent the observed energy consumption, *E*, and the solid line is the predicted energy consumption from Eq. (16).

For our algorithms, the rate of growth of problem size as a function of the number of vessels scaled as *α* + *ξN*_*V*_ , in which *α* and *ξ* are constants. In Big O notation, this may be expressed as O (*α* + *ξN*_*V*_). For small number of vessels (*N*_*V*_ ⪅ 10^4^), the first term is much larger than the second (*α*≫*ξN*_*V*_), so we saw an approximately constant computing time (Fig. 5), corresponding to high efficiency. For larger vascular networks, the second term predominated, and because *C* was approximately one, the observed efficiency was moderate. The findings are apparent by inspection of Eq. (9), its parameter values listed above, and Fig. 6.

Broadly, the CVDT codes exhibited strong and weak scalability across the entire range of vessels (6 ≤*N*_*V*_ ≤34 ×10^9^). Fig. 7(a) plots the measured speedup factor for strong scaling, which increased approximately sigmoidally (1 ≤ *S* ≤ 76) with increasing numbers of nodes (1 ≤ *N*_*c*_ ≤ 1024), asymptotically approaching the limit of 83 predicted by Amdahl’s Law in Eq. (12). Also for strong scaling, the fraction of the execution time *T* (1, *F*_0_), spent on tasks that could have been completed faster had they been computed in parallel, was estimated at *p*(*F*_0_) = 0.988 by fitting *S* values to Amdahl’s Law. Weak scaling tests reveals the parallel efficiency decreased monotonically (1 ≤*ε* ≤0.47) with increasing numbers of nodes (1 ≤*N*_*c*_ ≤1024) and approached an asymptotic value of 0.48 (Fig. 7(b)). The asymptote (*p*(*F*_0_ ×*N*_*c*_) = 0.48) corresponds to the fraction of *T* (1, *F*_0_ ×*N*_*c*_) that was spent on tasks that could have been reduced if executed on multiple compute nodes in Eq. (15). The findings regarding strong and weak scaling are consistent with Amdahl’s Law and Gustafson’s Law, respectively. Our results for strong and weak scalability are important because they provide the first evidence of the computational feasibility of a 34 billion vessel CVDT.

**Fig. 7.**
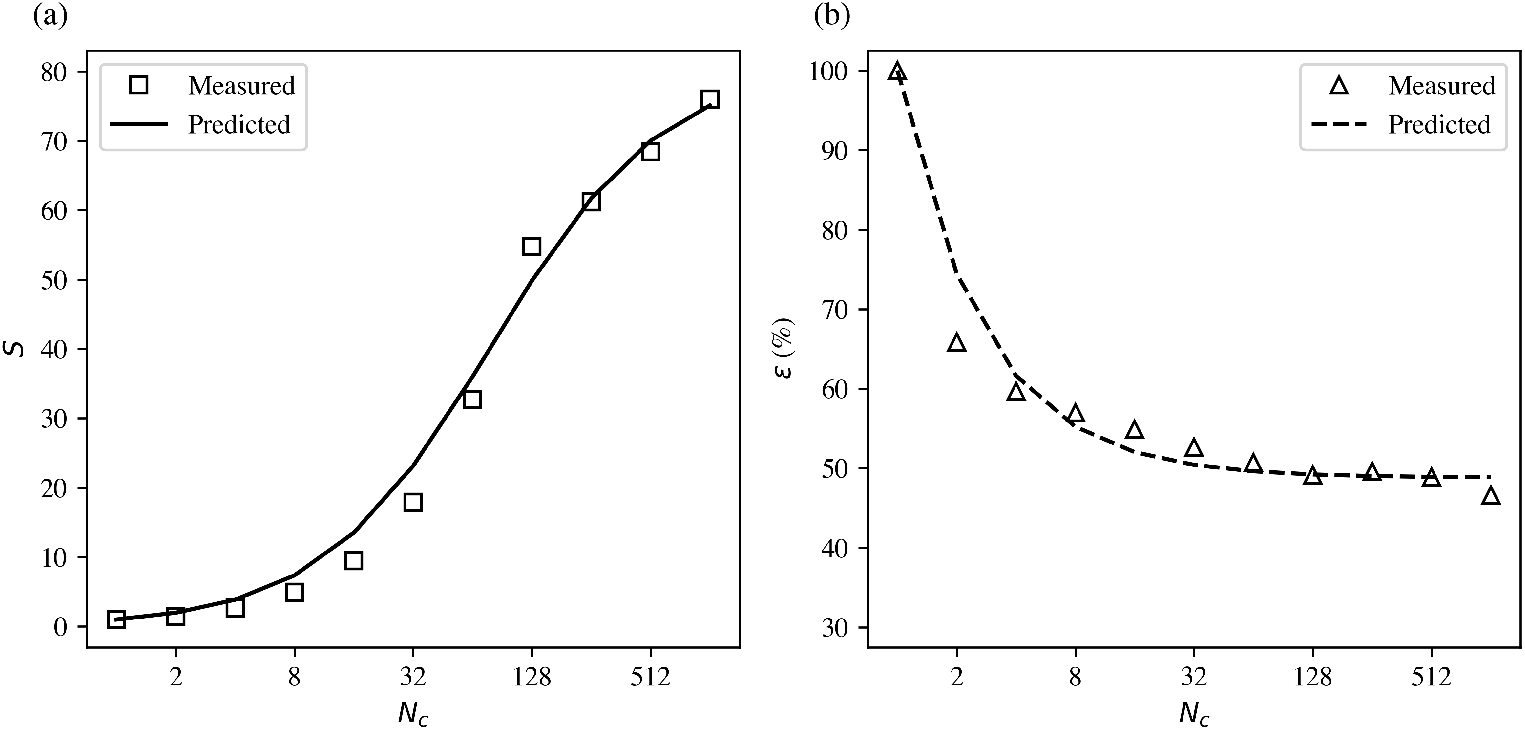
(a) Speedup, *S*, versus the number of compute nodes, *N*^*c*^ , measured for all algorithms combined (vascularization, partitioning & preprocessing, and blood flow calculations). The squares denote the measured speedup values from Eq. (10) and the solid line is an empirical fit of the speedup to Amdahl’s Law from Eq. (11). (b) Parallel efficiency, *ε*, versus number of compute nodes, *N*^*c*^ , for all algorithms combined. The triangles denote the measured parallel efficiency from Eq. (13) and the dashed line is an empirical fit of the parallel efficiency to Gustafson’s Law from Eq. (14).

Using Eq. (8), we estimated the minimum wall-clock time for floating point operations, *T*_fp_, between 2 and 34 generations. Fig. 6 displays the measured values of *T*_fp_ and those reported by Eq. (8). The maximum absolute difference between the measured and observed times was 43 ms with an average absolute difference of 7.2 ms. The maximum relative difference was 160% and the average difference was 40%. The root-mean-square difference was 1.6%. The differences were attributed to the Supercomputer Fugaku’s node allocation software, which throttles the CPU clock speed during periods of high traffic on the cluster. Fig. 6 shows the measured and estimated energy usage on Supercomputer Fugaku. The maximum relative difference was 31% and the average was 10%. The baseline power consumption value, *β*, was approximately 1.5 Wh on Supercomputer Fugaku.

### E. Validation

We validated the accuracy of the predicted blood pressures and flow rates by comparing to corresponding test results obtained using a previously validated commercial computational fluid dynamics simulation software. The maximum relative difference found was less than 1% for all vessels (24). Fig. 8 displays the volumetric blood flow rates in a 62-vessel network (5 generations) projected onto a 2-dimensional illustration. As expected, vessels in the same generation have equal blood flow rates, in which the arterial and venous trees are mirror symmetric.

**Fig. 8.**
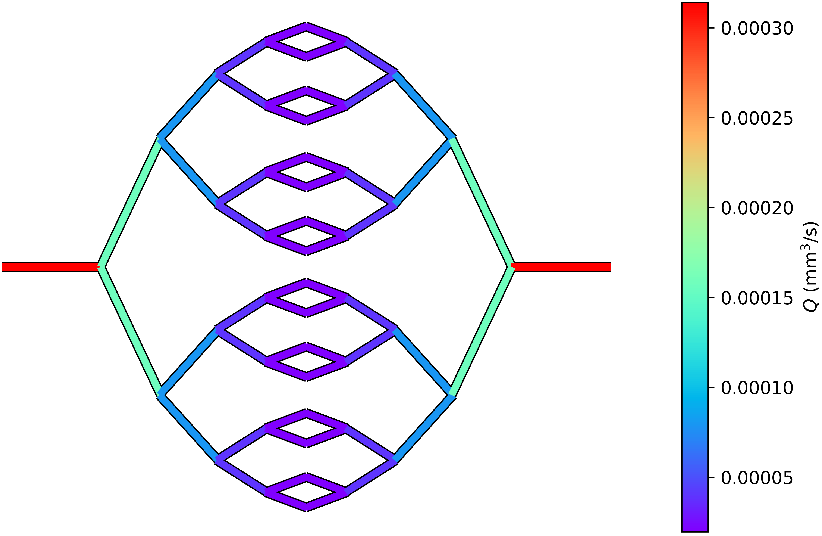
Illustrative example of calculated blood flow rates, *Q*, overlayed on a 2-dimensional 62-vessel network. For validation, boundary conditions for blood pressures and flow rate were set equal to those described by Donahue *et al*. (24).

To ascertain reproducibility, we performed regression tests by comparing our results to those found by three alternative methods (Section C). First, we compared our calculated blood flow rates (Section A.3) to those found by Eq. (17) and found that the relative difference in flow rates for each vessel was less than 0.05%. We also performed regression tests by comparing our results to the previous algorithms from our laboratory and found the maximum relative difference to be less than 0.05%. Finally, we implemented modified versions of our algorithms in the Python programming language for simulation of smaller network sizes on a personal computer with 8 GB of memory. The personal computer was capable of generating vasculature and solving blood flow rates in networks with up to 65, 534 vessels (14 generations) in 24.6 s. The maximum relative difference for blood flow rates was less than 0.5% for all vessels.

## Discussion

In this study, we developed methods to simulate blood flow through a vascular network containing 34 billion vessels, approximately the same number of vessels in the entire human body. The major finding of this study is that, for the first time, the feasibility of CVDT modeling was successfully demonstrated. The approach is computationally scalable with parallel computing systems up to tera-FLOP and tera-byte scales. Secondary findings shed new light on the portability, energy consumption and other performance of cardiovascular simulations.

The long-term implication of these findings is that this approach, namely, first-principles biophysical modeling of the cardiovascular system, is a viable pathway to realizing more realistic CVDTs for eventual use in personalized medicine. The simplified full-body model presented in this report proves key capabilities that will be needed for future CVDTs which that will find application in many areas of human health, including diagnosis and treatment of diseases, performance enhancement, trauma, and metabolism. The short-term implication of this study is that future cardiovascular researchers will soon have tools available to simulate blood circulation in the whole body. A first-principles, whole organism approach will open new avenues of inquiry, enable new approaches to hypothesis generation, and facilitate the design of experimental research studies, *e*.*g*., mechanistic and pre-clinical animal studies.

The findings of this study are generally coherent with the literature. In particular, the study is similar to that from Donahue *et al*. (24) in two important ways. Quantitatively, the present study achieved another significant increase in the vasculature size, *i*.*e*., a factor of two increase to 34 billion vessels. In addition, speed and scalability are now superior and better understood. Smaller, though no less impressive, increases in capacity were reported by Linninger *et al*., for detailed vascular networks of 256,000 vessels in 2013 (21) and 500,000 vessels in 2019 (22). The number of vessels and vascular density of our network was also consistent with those of the entire human body, as reported by Schmidt and Thews (12). Taken together with other recent literature, our results suggest that the frontier of computational vascular modeling is poised to expand rapidly and, in the process, open new avenues in large multi-scale biophysical modeling, including CVDTs for research and clinical applications.

Notable strengths of this study include extensive efforts to better understand computational requirements of CVDTs, which should facilitate future research. This includes computational requirements for arbitrarily-sized networks (*e*.*g*., time, scaling, energy consumption) and characteristics of the software (*e*.*g*., portability, ease of use, and extensibility). Another important strength of this study is the substantial efforts to demonstrate reproducibility, such as utilizing only non-proprietary or open-source software. Study methods were reproduced almost independent of those of Donahue *et al*. (24), increasing confidence in the approach and their general applicability. It is noteworthy that this is the first study to model a vascular network containing the number of vessels in the human body, steady-state blood flow, and scaling laws for CVDT simulations. These are major milestones on the path to full-fledged CVDTs.

The study has several limitations, including simplified vessel geometry, steady state blood flow, and other simplifications that were necessary to prove computational feasibility at scale using currently-available hardware. These limitations are also examples of general characteristics of computational CV research at scale. Specifically, the task of simulating the entire CV system with full biophysical realism vastly exceeds current computing capabilities. Therefore, all studies must limit realism, problem size, or both to achieve computational feasibility. In other words, the main purpose of research in this area will be to selectively increase biophysical realism while maintaining computational feasibility. Furthermore, simplified first-principles modeling methods, such as ours, are well suited for future refinement, *e*.*g*., as additional computing power becomes available.

To overcome several specific limitations of this study, studies are underway in our laboratory to explore scalability of time-dependent blood flow, vasodilation, diffusive oxygen transport from blood to tissue, and tracing of objects in the blood. Other limitations can be overcome by utilizing models from other laboratories. For example, a few other groups have already reported the computational requirements to simulate more anatomically detailed models of comparatively small numbers of vessels (9, 22). Other laboratories are utilizing magnetic resonance imaging (MRI) data from human subjects to produce highly realistic vascular geometry models. A study aimed at ascertaining feasibility and scalability of CVDTs with higher biophysical realism is currently underway in our laboratory. For all of these reasons, the use in the present study of a simplified vascular network is not a major limitation.

Clearly, the primitive CVDT presented here underscores that this area of research is in its infancy. It also appears that, for the foreseeable future, all such CVDTs will need to strike a judicious balance between speed of execution and the amount of biophysical realism that is modeled. Priorities for additional modeling realism include the incorporation of time-dependent blood flow, elasticity of blood vessels, vasoconstriction, vessel rupture, and oxygen saturation. Because these additions will require at least peta-scale computing resources, additional research on the computational scalability of CVDTs will be essential.

Our laboratory is interested in utilizing CDVTs to improve outcomes for cancer patients. With approximately 90% of all cancer patients dying from metastases (65), research is urgently needed to develop novel strategies to prevent the spread of primary tumors via circulating tumor cells. Specifically, this will require modeling intravasation of tumor cells through vessel walls into the bloodstream, transport of tumor cells throughout the circulatory system, and extravasation of these cells through vascular walls into healthy tissues. Similarly, CVDTs may help to understand or even prevent side effects of cancer radiotherapy. Radiation injury of blood vessels can lead to reduced blood flow (66–68), which reduces the rates of perfusion and diffusion of oxygen (69). Hypoxia can lead to tumors becoming resistant to radiation and can also induce necrosis. Reducing or preventing radioresistance and radiation necrosis has the potential to improve outcomes for many cancer patients.

## Conclusions

For the first time, this study demonstrated that it is computationally feasibility to simulate blood flow through a vascular network comprising 34 billion vessels, the number in the human body. In addition, the results revealed good computational scalability, which indicates that this type of computational modeling is well suited to solution with tera-scale parallel supercomputing techniques. These findings suggest that it will be feasible to continue to enhance the realism of existing biophysical models (*e*.*g*., pulsatile flow and more realistic vessel geometries) and add additional ones (vessel changes due to radiation injury and vaso-constriction and -dilation). The findings also suggest the feasibility to further develop cardiovascular simulation techniques to the point where they are capable to impact research on important cardiovascular health challenges, such as stroke, radiation injury, and circulating tumor cells. The general approach appears to be well suited for eventual utilization in personalized medicine, including in cardiovascular digital twins.

## Acknowledgments

The authors would like to thank the Bella Bowman Foundation for financial support in part for this research project. We acknowledge Stony Brook Research Computing and Cyberinfrastructure Group and the Institute for Advanced Computational Science at Stony Brook University for access to the high-performance Ookami computing system, which was made possible by a grant from the National Science Foundation (1927880). This work used computational resources of the Supercomputer Fugaku provided by RIKEN through the HPCI System Research Project (Project ID: hp210311). We are grateful to the support staff at LSU’s Center for Computation and Technology for their generous technical assistance. Support was also provided via research grants from NASA 80NSSC21K0544 and the National Science Foundation (2229751 and 2229751). This work was also supported by the U.S. Department of Energy through the Los Alamos National Laboratory. Los Alamos National Laboratory is operated by Triad National Security, LLC, for the National Nuclear Security Administration of U.S. Department of Energy (Contract No. 89233218CNA000001) LA-UR-24-30400.

